# Time-Dependent Increase in Susceptibility and Severity of Secondary Bacterial Infection during SARS-CoV-2 Infection

**DOI:** 10.1101/2022.02.28.482305

**Authors:** Amanda P. Smith, Evan P. Williams, Taylor R. Plunkett, Muneeswaran Selvaraj, Lindey C. Lane, Lillian Zalduondo, Yi Xue, Peter Vogel, Rudragouda Channappanavar, Colleen B. Jonsson, Amber M. Smith

## Abstract

Secondary bacterial infections can exacerbate SARS-CoV-2 infection, but their prevalence and impact remain poorly understood. Here, we established that a mild to moderate SARS-CoV-2 infection increased the risk of pneumococcal coinfection in a time-dependent, but sex-independent, manner in the transgenic K18-hACE mouse model of COVID-19. Bacterial coinfection was not established at 3 d post-virus, but increased lethality was observed when the bacteria was initiated at 5 or 7 d post-virus infection (pvi). Bacterial outgrowth was accompanied by neutrophilia in the groups coinfected at 7 d pvi and reductions in B cells, T cells, IL-6, IL-15, IL-18, and LIF were present in groups coinfected at 5 d pvi. However, viral burden, lung pathology, cytokines, chemokines, and immune cell activation were largely unchanged after bacterial coinfection. Examining surviving animals more than a week after infection resolution suggested that immune cell activation remained high and was exacerbated in the lungs of coinfected animals compared with SARS-CoV-2 infection alone. These data suggest that SARS-CoV-2 increases susceptibility and pathogenicity to bacterial coinfection, and further studies are needed to understand and combat disease associated with bacterial pneumonia in COVID-19 patients.

## Introduction

Throughout the coronavirus disease 19 (COVID-19) pandemic caused by the severe acute respiratory syndrome coronavirus 2 (SARS-CoV-2), there have been case reports, multi-center cohort studies, systematic reviews, and meta-analyses assessing the extent and severity of coinfections with secondary pathogens including viruses, fungi, and bacteria^1–31^. Although coinfection rates varied across studies, some studies suggested that coinfecting respiratory bacteria were predictors of severe SARS-CoV-2-related disease and mortality^23–31^. Bacterial pathogens that were detected included *Mycoplasma pneumoniae, Legionella pneumophila, Chlamydophila pneumoniae, Klebsiella pneumoniae, Pseudomonas aeruginosa, Haemophilus influenzae, Acinetobacter baumanii, Staphylococcus aureus,* and *Streptococcus pneumoniae* (pneumococcus). Pneumococcus, which is a major cause of community-acquired pneumonia^32–34^, was detected by throat swab in 0.8%^8^ to 7.2%^5^ of hospitalized COVID-19 patients not requiring intensive care unit (ICU) admission or invasive respiratory support, while the frequency tended to be higher (6.5%^24^ to 59.5%^4^) in patients with severe respiratory distress. Because bacterial transmission has largely been dampened by non-pharmaceutical measures (e.g., masking and physical distancing), it is important to understand whether SARS-CoV-2 infection predisposes individuals to bacterial infections and, if so, what clinical and immunological changes occur as a result of coinfection.

In general, viral-bacterial coinfections are not uncommon, where *S. aureus* and pneumococcus are widely documented as complicating pathogens during infection with other viruses, most notably influenza A virus (IAV)^Reviewed in 35–46^. During influenza pandemics, 45-95% of the mortality has been attributed to bacterial coinfections^47–50^. Fortunately, the impact of these complications has appeared to be lower during the SARS-CoV-2 pandemic, but these could increase as novel variants arise and as SARS-CoV-2 becomes endemic. IAV and SARS-CoV-2 both cause infections that range from asymptomatic to severe, but SARS-CoV-2 has a longer incubation period, longer and more varied duration of viral shedding and symptoms, and more pathological effects on tissues outside of the respiratory tract^Reviewed in 51–54^. Although viral burden does not directly correlate to disease^55–61^, both viruses can induce significant lung damage^Reviewed in 52–54^. Some host responses also differ in timing and magnitude, including the delayed type I interferon (IFN-α,β), increased proinflammatory cytokines like TNF-a and IL-6, and reduced immune regulation that have been detected in COVID-19 patients^62–66^. Further, neutrophils and macrophages, which are important for efficient bacterial clearance during viral-bacterial coinfection^67–72^, are dysregulated during COVID-19^73–75^. Thus, the potential for bacterial invasion during SARS-CoV-2 infection may also differ from that observed in influenza infection with respect to timing and host-pathogen mechanisms.

While the investigation of viral and immune dynamics in the lower respiratory tract is difficult to assess in humans, they have been clarified in animal models. One study using SARS-CoV-1 suggested that bacteria can enhance pathogenicity of coronaviruses^76^, and numerous studies of influenza-bacterial coinfection indicate that susceptibility and pathogenicity of bacterial coinfections are time-dependent with the greatest mortality observed when bacteria is initiated at 7 d pvi^77^. The progressive increase in susceptibility to bacterial coinfection during influenza is largely due to the depletion and/or dysfunction of resident alveolar macrophages (AMΦ) during IAV infection, which is dynamic throughout the infection^55,67^ and maximal at 7 d pvi^55,67–69^. Following bacterial establishment, dysfunction of neutrophils^78–81^, which may be in part facilitated by bacterial metabolic interactions^82^ and type I IFNs^71,82,83^, and additional depletion of AMΦ^55^ contribute to bacterial growth and coinfection pathogenesis^Reviewed in 39–41,45,84,85^. Currently, the effect of SARS-CoV-2 infection on AMΦs remains somewhat unclear, although human, murine, and *in vitro* data indicate that AMΦs become productively infected with SARS-CoV-2, leading to altered cytokine production and responsiveness^86–89^. In addition, SARS-CoV-2 seems particularly adept at delaying and avoiding innate immune responses, resulting in delayed or decreased T cell responses, accumulation of neutrophils and inflammatory monocytes, and enhanced lung pathology^Reviewed in 90–93^. IAV also has mechanisms of immune evasion^Reviewed in 94,95^ but induces a robust CD8^+^ T cell response in the lungs that efficiently clears virus. During IAV-pneumococcal coinfection, CD8^+^ T cells are depleted^96^, and viral loads rebound^55,68,82^. Mechanisms for both of these are being investigated, but direct viral-bacterial interactions^97^ that allow the virus to enter new areas of the lung in addition to a bacterial-mediated increase in virus production^55,68,98^ contribute to the increased viral loads. However, these effects are overshadowed by the robust bacterial growth and bacterial-mediated effects on host responses. Given these potential mechanisms and the reported myeloid dysfunction^73–75^, delayed IFN responses^62–66^, and CD8^+^ T cell depletion^99–103^ during SARS-CoV-2, a better understanding of the potential for bacterial invasion and the effects of coinfection on immune cell, viral, and pathological dynamics is needed and the focus of this study. To assess bacterial susceptibility during COVID-19 and determine whether a synergism exists between SARS-CoV-2 and pneumococcus, we infected K18-hACE2 mice with a low dose of SARS-CoV-2 to initiate a mild-moderate infection and coinfected the animals 3, 5, or 7 days later with pneumococcus. Bacteria were unable to establish at 3 d post-virus infection (pvi), but coinfections at 5 or 7 d pvi resulted in increased lethality in a sex-independent manner. Although viral dynamics and lung pathology were unchanged within the first 24 h of coinfection, select immune cells and proinflammatory cytokines were decreased in the lungs of animals coinfected at 5 d pvi but not at 7 d pvi. These findings support the increased susceptibility of SARS-CoV-2-infected individuals to bacteria and highlight numerous distinct features from other viral-bacterial coinfections.

## Results

### Time-dependent increases in lethality during SARS-CoV-2-pneumococcal coinfection

To examine the susceptibility and pathogenicity of pneumococcus coinfection during SARS-CoV-2 infection, K18-hACE2 mice (male and female, 10 to 13 weeks old) were infected with 250 PFU of SARS-CoV-2 or PBS followed by 10^3^ CFU of pneumococcal strain D39 (coinfected) or PBS (mock coinfected) at either 3, 5, or 7 d pvi. During mock coinfection, the selected viral dose was lethal in 35% of mice (**Figure 1**A) and caused weight loss from 5 to 11 d pvi with maximum weight loss (average 7%) at 8 d pvi (**Figure 1**B) and clinical scores peaking at 6 d pvi (**Figure 1**C). In the absence of viral infection, the selected bacterial dose was lethal in 1/6 mice (17% lethality) at 4 d post bacterial infection (pbi) (Fig S1A) and caused only mild, transient weight loss (~3%) (Fig S1B) and increased temperatures (Fig S1C) after 1 to 2 d pbi.

**Figure 1:**
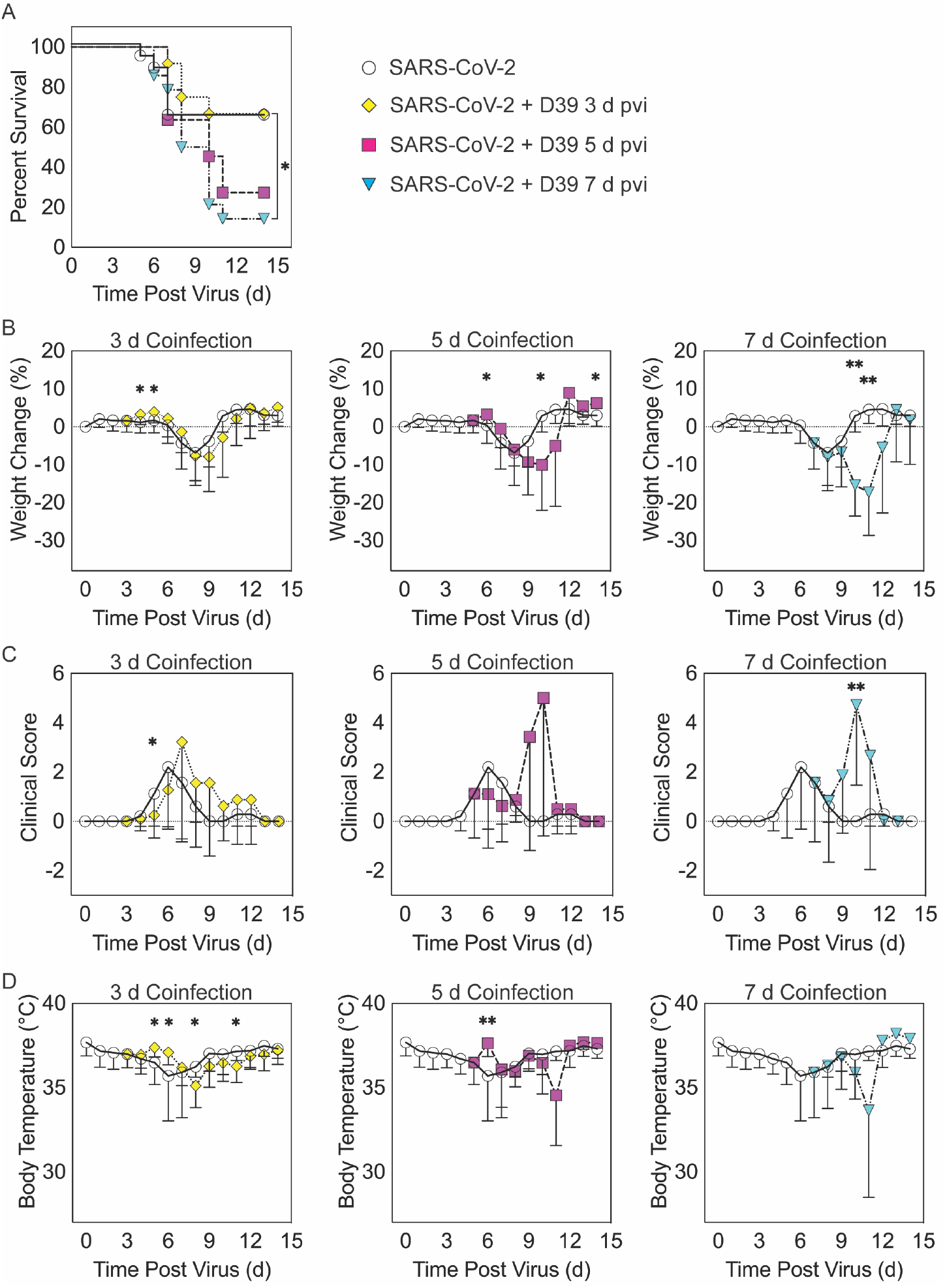
SARS-CoV-2-pneumococcal coinfection in K18-hACE2 mice. Kaplan-Meier survival curves (A), percent weight loss (B), cumulative clinical score (C), and temperature (D) of mice infected with SARS-CoV-2 (250 PFU; white circles, solid lines) followed by infection with 10^3^ CFU D39 at 3 d (yellow diamonds, dotted lines), 5 d (magenta squares, dashed lines), or 7 d (cyan triangles, dash-dotted lines) pvi. Data are shown as the mean ± standard deviation (SD) and significant differences are indicated by *, *P* < 0.05; **, *P* < 0.01 for comparisons between SARS-CoV-2 infection and SARS-CoV-2-pneumococcal coinfection.

When the bacterial coinfection was initiated at 3 d pvi, lethality was not enhanced (*P* = 0.73) (Figure 1A). Interestingly, weight loss in coinfected animals was reduced at 1 d (*P* = 0.03) and 2 d (*P* = 0.04) pbi (Figure 1B) and the cumulative clinical score was lower at 2 d pbi (*P* = 0.03) (Figure 1C) compared with mock coinfected controls. In addition, the temperature of coinfected animals was higher at 2 d (*P* = 0.003) and 3 d (*P* = 0.01) pbi and lower at 5 d (*P* = 0.02) and 8 d (*P* = 0.045) pbi (Figure 1D). A coinfection initiated at 5 d pvi was slightly more lethal than the SARS-CoV-2 infection alone, where additional mortality was observed at 5 to 6 d pbi, but this was not statistically significant (*P* = 0.14) (Figure 1A). The average weight loss was reduced (*P* = 0.01) and temperature was increased (*P* = 0.001) at 1 d pbi in the coinfected animals (Figure 1B and D). Coinfected animals lost more weight than animals infected with SARS-CoV-2 alone at 5 d pbi (*P* = 0.03) (Figure 1B), but no significant difference in their clinical scores was detected (Figure 1C). Comparatively, a coinfection at 7 d pvi was significantly more severe than SARS-CoV-2 infection alone (*P* = 0.03) and resulted in additional lethality at earlier times than the coinfection at 5 d pvi, with additional animals succumbing to the infection within 1, 3, or 4 d pbi (Figure 1A). Significantly more weight loss at 3 d (*P* < 0.001) and 4 d (*P* = 0.002) pbi (Figure 1B) and higher clinical scores at 3 d pbi (*P* = 0.01) (Figure 1C) occurred without altering temperature (Figure 1D).

### SARS-CoV-2 coinfection increased bacterial loads but not viral loads

To evaluate whether SARS-CoV-2-bacterial coinfection alters pathogen burden, we measured viral loads in the lung and bacterial loads in the lung and blood of infected animals. In mice infected with bacteria alone or with SARS-CoV-2 followed by bacteria at 3 d pvi, no bacteria were recovered from the lungs of 7/8 mice at 24 h pbi (Figure 2A and Figure S1D). However, when the bacteria was introduced at 5 d pvi, bacterial loads in the lung remained at a level similar to the inoculum in 7/8 mice and was cleared in 1/8 mice (Figure 2A). Bacteria were not detected in the blood of mice infected with bacteria alone (data not shown) or SARS-CoV-2-bacteria coinfected at 3 or 5 d pvi (Figure 2B). However, in mice coinfected at 7 d pvi, significant bacterial growth occurred in the lungs of all animals and the blood of some animals (3/7) with titers reaching 4.4 to 7.9 log™ CFU/lung (Figure 2A) and 4.1 to 6.6 log_10_ CFU/mL (Figure 2B), respectively, within 24 h pbi.

**Figure 2:**
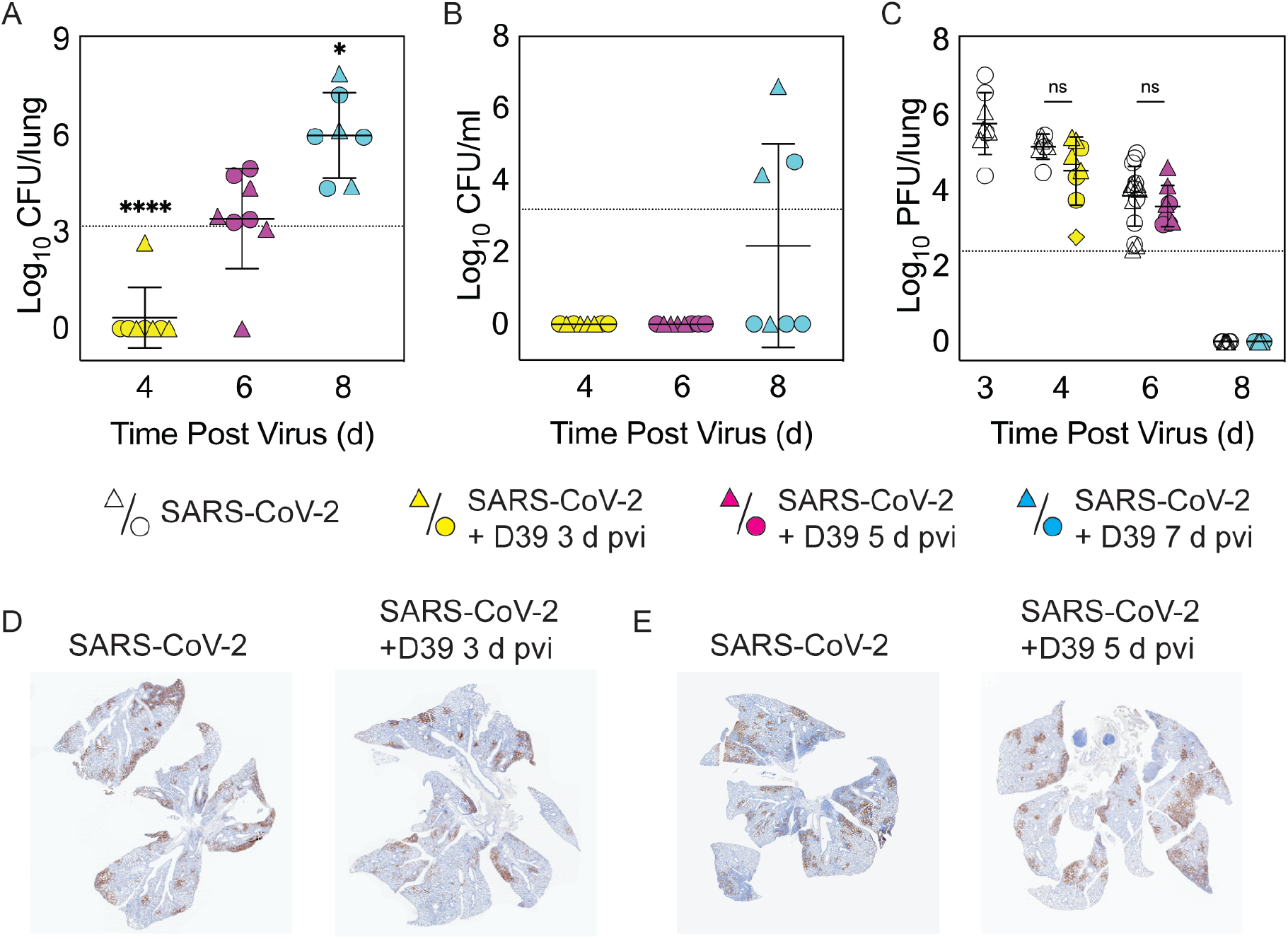
Dynamics of pathogen loads during SARS-CoV-2 infection and pneumococcal coinfection. Lung bacterial loads (CFU/lung) (A), blood bacterial loads (B), and lung viral loads (PFU/lung) (C) in female (circles) and male (triangles) mice infected with SARS-CoV-2 (250 PFU; white) followed by infection with 10^3^ CFU D39 at 3 d (yellow), 5 d (magenta), or 7 d (cyan) pvi. Each symbol represents a single mouse and the mean ± standard deviation (SD) are for combined male and female groups. Significant differences are indicated by ns, not significant; *, *P* < 0.05; ***, *P* < 0.0001. For bacterial titers, comparison was with the inoculum (dotted line). (D-E) Representative immunohistochemical staining of SARS-CoV-2 nucleocapsid protein in the lungs of mice 24 h after they were infected with SARS-CoV-2 (250 PFU) ± 10^3^ CFU D39 at 3 d (D) or at 5 d (E) pvi.

Pulmonary viral loads were unchanged by bacterial coinfection whether coinfection was initiated at 3 d (*P* = 0.12) or 5 d (*P* = 0.18) pvi (Figure 2C) and the amount and distribution of viral antigen in the lung tissue were also unchanged (Figure 2D and E). The virus was cleared by 8 d pvi in the groups that were either mock coinfected or bacterial coinfected at 7 d pvi (Figure 2C). No significant differences were found in viral or bacterial loads between males and females.

### Select changes in pulmonary immune responses after SARS-CoV-2-pneumococcal coinfection

To investigate whether bacterial coinfection altered immune response dynamics, several immune cells, cytokines, and chemokines were quantified in the lung 24 h after mock coinfection or bacterial coinfection in SARS-CoV-2 infected mice (Figure 3 and 4, Figure S3 to S6). In animals infected with SARS-CoV-2 only, natural killer (NK) T cells (Figure S3D) and total CD19^+^ B cells (Figure 3E) were reduced at 4 d pvi compared with naïve (*P* = 0.007 and *P* = 0.018, respectively). The absolute numbers of other cells were unchanged at this time point (Figure 3 and Figure S3); however, increases in the proportion of activated (CD69^+^) immune cells were evident (Figure S4). SARS-CoV-2 infection also resulted in many cytokines and chemokines above baseline levels (all *P* < 0.05) throughout the infection, including IFN-γ, IL-1β, IL-4, IL-28, CXCL10, GM-CSF, LIF, CCL2, CCL7, MIP-1α, MIP-1β, RANTES, IFN-α, and IFN-β. IL-5, IL-6, IL-15, IL-18, M-CSF, and TNF-α were elevated at both 4 d and 6 d pvi while CXCL5, CXCL1, G-CSF, IL-3, IL-13, and IL-17A were increased only at 6 d pvi. MIP-2α, IL-2, and IL-22 were elevated at 6 d and 10 d pvi, and increased IL-10 and IL-23 were detected only at 8 d pvi (absolute values of cytokines are in Figure 4 and Figure S5; log2 changes over naïve in Figure S6).

**Figure 3:**
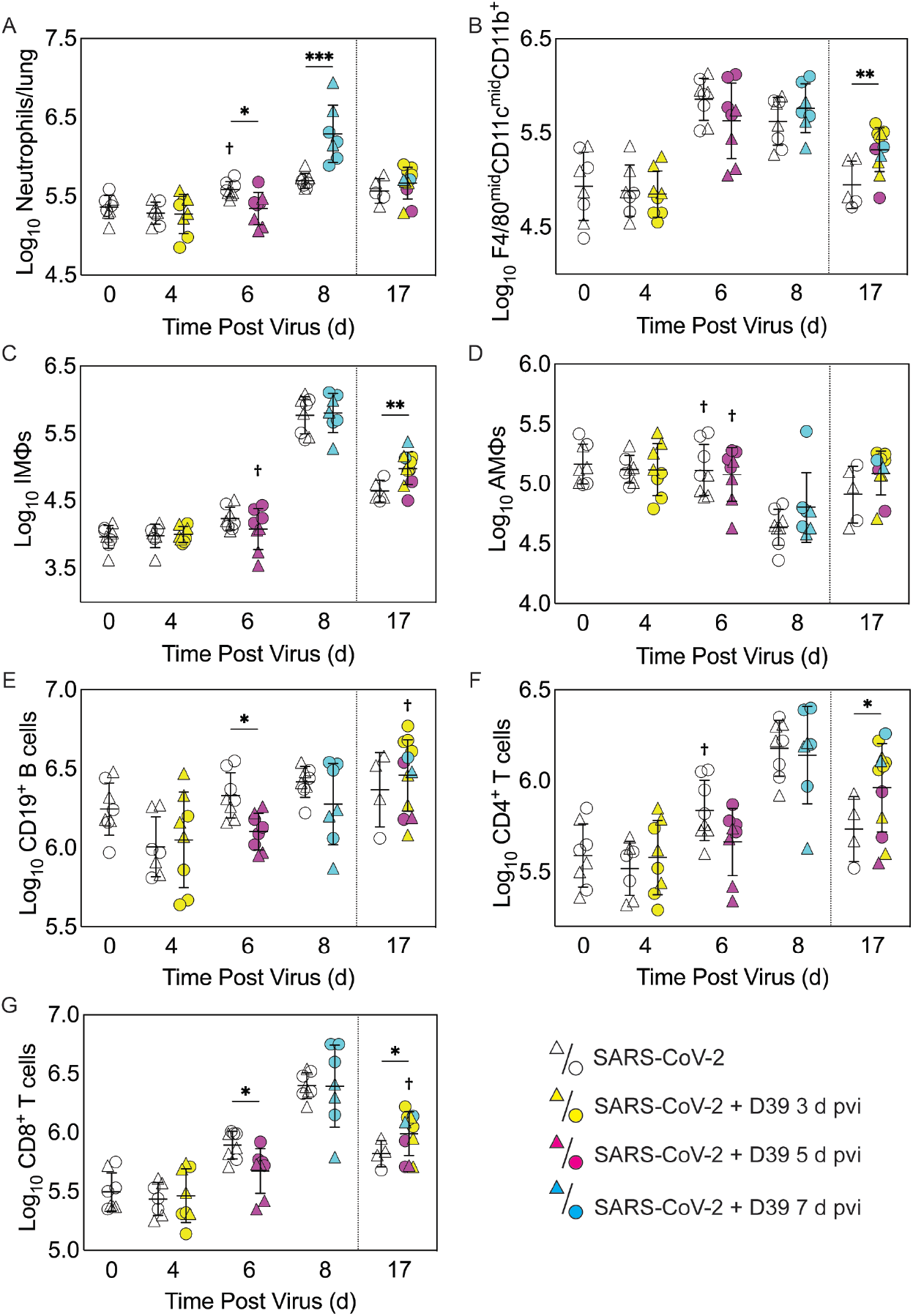
Immune cell dynamics during SARS-CoV-2 infection and pneumococcal coinfection. Total neutrophils (A), F4/80^mid^CD11c^mid^CD11b^+^ monocytes/macrophages (B), inflammatory macrophages (iMΦ) (F4/80^hi^CD11c^hi^CD11b^+^) (C), alveolar macrophages (AMΦ) (F4/80^hi^CD11c^hi^CD11b^-^MHC-II^low/-^) (D), CD19^+^ B cells (E), CD4^+^ T cells (F), and CD8^+^ T cells (G) in the lungs of female (circles) and male (triangles) mice infected with SARS-CoV-2 (250 PFU; open symbols) followed by infection with 10^3^ CFU D39 at 3 d (yellow), 5 d (magenta), or 7 d (cyan) pvi. Each symbol represents a single mouse and the mean ± standard deviation (SD) are for combined male and female groups. Significant differences are indicated by *, *P* < 0.05; **, *P* < 0.01; ***, *P* < 0.001 for comparisons between indicated groups and by †, *P* < 0.05 for differences between males and females within a group or between coinfection times within 17 d group.

**Figure 4:**
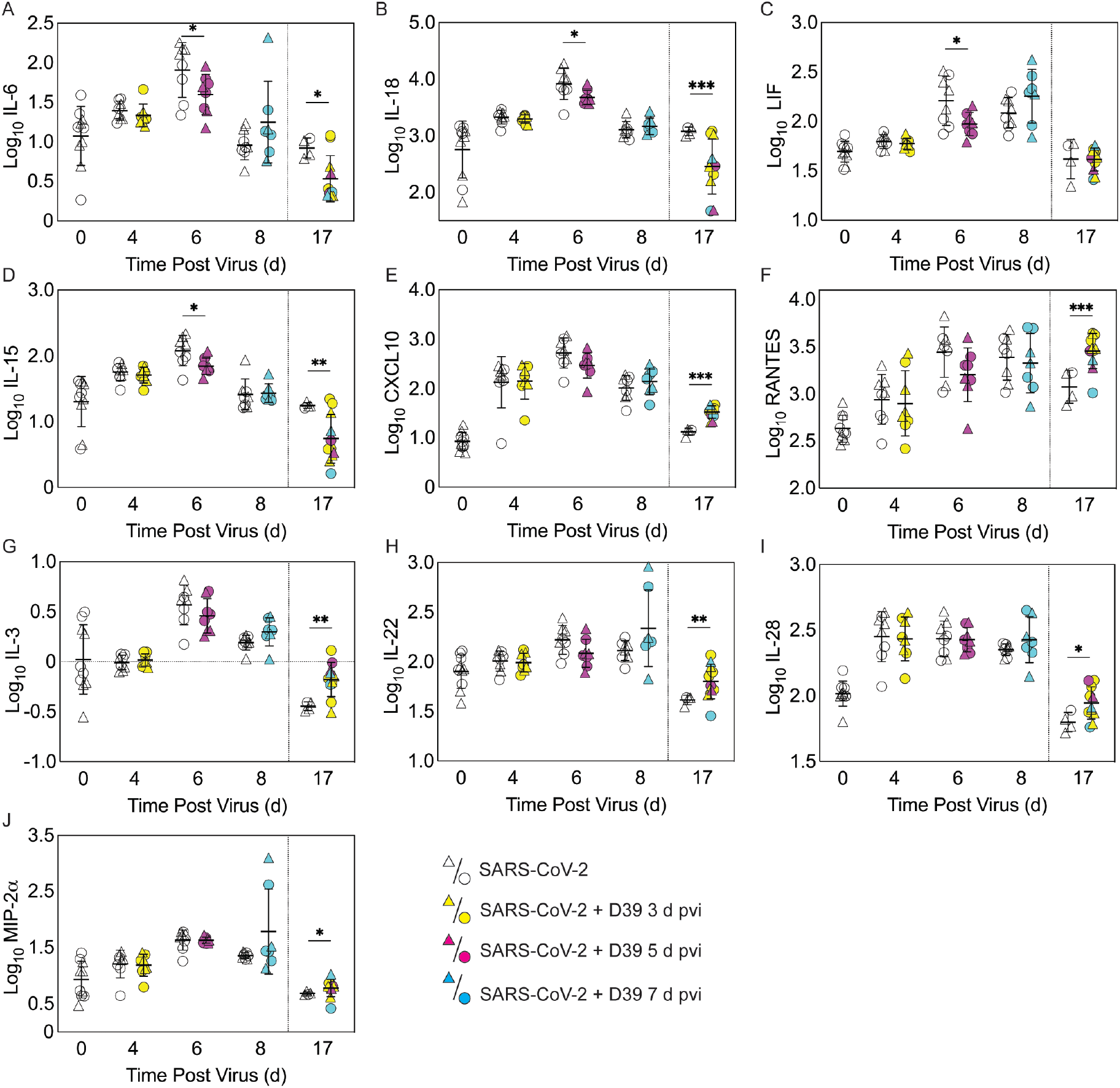
Pulmonary cytokines and chemokines during SARS-CoV-2 infection and SARS-CoV-2-pneumococcal coinfection. Total IL-6 (A), IL-18 (B), LIF (C), IL-15 (D), CXCL10 (E), RANTES (F), IL-3 (G), IL-22 (H), IL-28 (I), and MIP-2α (J) in the lungs of female (circles) and male (triangle) mice infected with SARS-CoV-2 (250 PFU; white) followed by infection with 10^3^ CFU D39 at 3 d (yellow), 5 d (magenta), or 7 d (cyan) pvi. Each symbol represents a single mouse and the mean ± standard deviation (SD) are for combined male and female groups. Significant differences are indicated by *, *P* < 0.05; **, *P* < 0.01; ***, *P* < 0.001 for comparisons between indicated groups. Plots depicting additional cytokine and chemokine quantities (absolute log_10_-10 picograms) are in Figure S5 and a heatmap representing the normalized quantity (average log2 change over naïve) is in Figures S6.

As expected, a significant influx of CD45^+^ immune cells was evident at 6 and 8 d pvi in animals infected with SARS-CoV-2 only (both *P* < 0.001) (Figure S3A), including neutrophils (Ly6G^hi^; both *P* < 0.01; Figure 3A), the F4/80^mid^CD11c^mid^CD11b^+^ monocyte/macrophage subset (both *P* < 0.001; Figure 3B), inflammatory macrophages (F4/80^hi^CD11c^hi^CD11b^+^, iMΦ; *P* = 0.02 and *P* < 0.001, respectively; Figure 3C), F4/80^mid^CD11c^-^ cells (both *P* < 0.001; Figure S3B), NK cells (both *P* < 0.001; Figure S3C), CD4^+^ T cells (*P* = 0.02 and *P* < 0.001, respectively; Figure 3F), and CD8^+^ T cells (both *P* < 0.001; Figure 3G). Unlike the pathogen loads, some of the immune cells were different between males and female that were mock coinfected at 5 d pvi, including neutrophils (*P* = 0.047), resident alveolar macrophages (F4/80^hi^CD11c^hi^CD11b^-^MHC-II^low/-^, AMΦ; *P* = 0.047), CD4^+^ T cells (*P* = 0.02), NK cells (*P* = 0.03), and NK T cells (*P* = 0.02), which were higher in females than males.

In the groups coinfected with bacteria at 3 d pvi, no changes were observed in the absolute number (Figure 3 and Figure S3) or activation (Figure S4) of any quantified immune cell subset or the amount of cytokines and cytokines (Figure 4 and Figure S5) within 24 h pbi compared with mock coinfection. A bacterial coinfection at 5 d pvi resulted in fewer total CD45+ cells (*P* = 0.03; Fig S3A), including neutrophils (Figure 3A), CD19^+^ B cells (Figure 3E), CD8^+^ T cells (Figure 3G), and F4/80^mid^CD11c^-^ cells (Fig S3B) (all *P* < 0.05) compared with the mock coinfected groups. In addition, iMΦ (*P* = 0.01) and AMΦ (*P* = 0.047) were again higher in females than males following coinfection at 5 d pvi (Figure 3C and D). The extent of activation was not different between the mock coinfection and bacterial coinfection at 5 d pvi (Figure S4), but reduced IL-6, IL-18, LIF (all P = 0.04), and IL-15 (P = 0.02) was observed at 24 h pbi (Figure 4A-D).

Coinfection at 7 d pvi induced a significant increase in neutrophils at 24 h pbi (P < 0.001) (Figure 3A) without altering the number or activation of any other immune cell quantified (Figure 3, Figure S3 and S4). AMΦ were reduced in the mock coinfected group compared with naïve animals (P = 0.001) but were not different between the mock coinfection and bacterial coinfection (P = 0.29) (Figure 3D). Absolute cell numbers and activation did not differ between male and female mice following coinfection at 7 d pvi (Figure 3, Figure S3 and S4). Perhaps unexpectedly, none of the measured cytokines were significantly different between animals that were mock coinfected and animals that were bacterial coinfected at 7 d pvi (Figure 4 and Figure S5).

### Pneumococcal coinfection resulted in sustained increases in pulmonary immune responses after recovery

To investigate whether bacterial coinfection altered immune cell dynamics and activation in recovered animals, pulmonary immune cells, cytokines, and chemokines were quantified at 17 d pvi following mock coinfection or bacterial coinfection at 3, 5, or 7 d pvi. The number of iMΦ (*P* = 0.01) (Figure 3C) and CD8^+^ T cells (*P* = 0.02) (Figure 3G), as well as the activated proportion of iMΦ (*P* = 0.004), CD8^+^ T cells (*P* = 0.001), CD4^+^ T cells (*P* = 0.001), and CD19+ B cells (*P* = 0.005) (Figure S4), remained increased above naïve levels in the lungs of animals that recovered from SARS-CoV-2 infection alone. These changes were accompanied by elevated IFN-γ, CXCL10, and RANTES (*P* = 0.01, *P* = 0.03, and *P* = 0.04, respectively) at 17 d pvi compared to naïve (Figure 4, Figure S5 and S6). However, many measured cytokines and chemokines were below naive levels at 17 d pvi in the lungs of animals infected with SARS-CoV-2 only, including eotaxin, IL-2, IL-3, IL-17A, IL-22, IL-27, IL-28, M-CSF, and MIP-2α (all *P* < 0.05) (Figure 4, Figure S5 and S6).

A sustained increase in immune cell accumulation and activation was evident in animals that recovered from SARS-CoV-2-pneumococcal coinfection. At 17 d pvi, an increased absolute number and activated proportion of F4/80^mid^CD11 c^mid^CD11b^+^ monocytes/macrophages (*P* = 0.01; Figure 3B and Figure S4B), iMΦ (*P* = 0.01; Figure 3C and Figure S4C), and CD4^+^ and CD8^+^ T cells (*P* = 0.03 and 0.02, respectively; Figure 3F and G and Figure S4F and G) were present in coinfected mice compared with mock coinfected mice. Comparison between the coinfected groups indicated that more CD8^+^ T cells were present at 17 d pvi in mice that were coinfected at 3 d or 7 d pvi than those coinfected at 5 d pvi (both *P* = 0.02; Figure 3G). In addition, animals that recovered from a coinfection at 7 d pvi had more activated neutrophils or iMΦ than those who recovered from a coinfection at 3 d pvi (*P* = 0.04) or 5 d pvi (*P* = 0.03), respectively (Figure S4A and C). These changes were accompanied by higher levels of CXCL-10 (*P* < 0.001), MIP-2a (*P* = 0.04), IL-3 (*P* = 0.001), IL-22 (*P* < 0.008), IL-28 (*P* = 0.01), and RANTES (*P* < 0.001) in the lungs of mice that had recovered from a bacterial coinfection compared with those recovered from SARS-CoV-2 alone (17 d pvi; Figure 4E to J). In addition, G-CSF, CXCL-1, IL-1α, IL-6, IL-9, IL-10, IL-13, IL-15, IL-18, and TNF-α (all *P* < 0.05) were lower in coinfected animals than mock coinfected controls at 17 d pvi (Figure S5).

### Bacterial coinfection did not enhance lung pathology

To examine whether lung pathology was enhanced during SARS-CoV-2-pneumococcal coinfection, we assessed seven pathological features (endothelial hypertrophy/margination, peribronchiolar/perivascular lymphoid cells, interstitial inflammation/septal thickening, alveolar inflammation, alveolar edema/hemorrhage, the extent of alveolar involvement, and consolidation (Figure 5). There were no significant differences in any of these measurements between mock coinfected animals and those coinfected with bacteria at 3 or 5 d pvi at either 24 h pbi or 17 d pvi.

**Figure 5:**
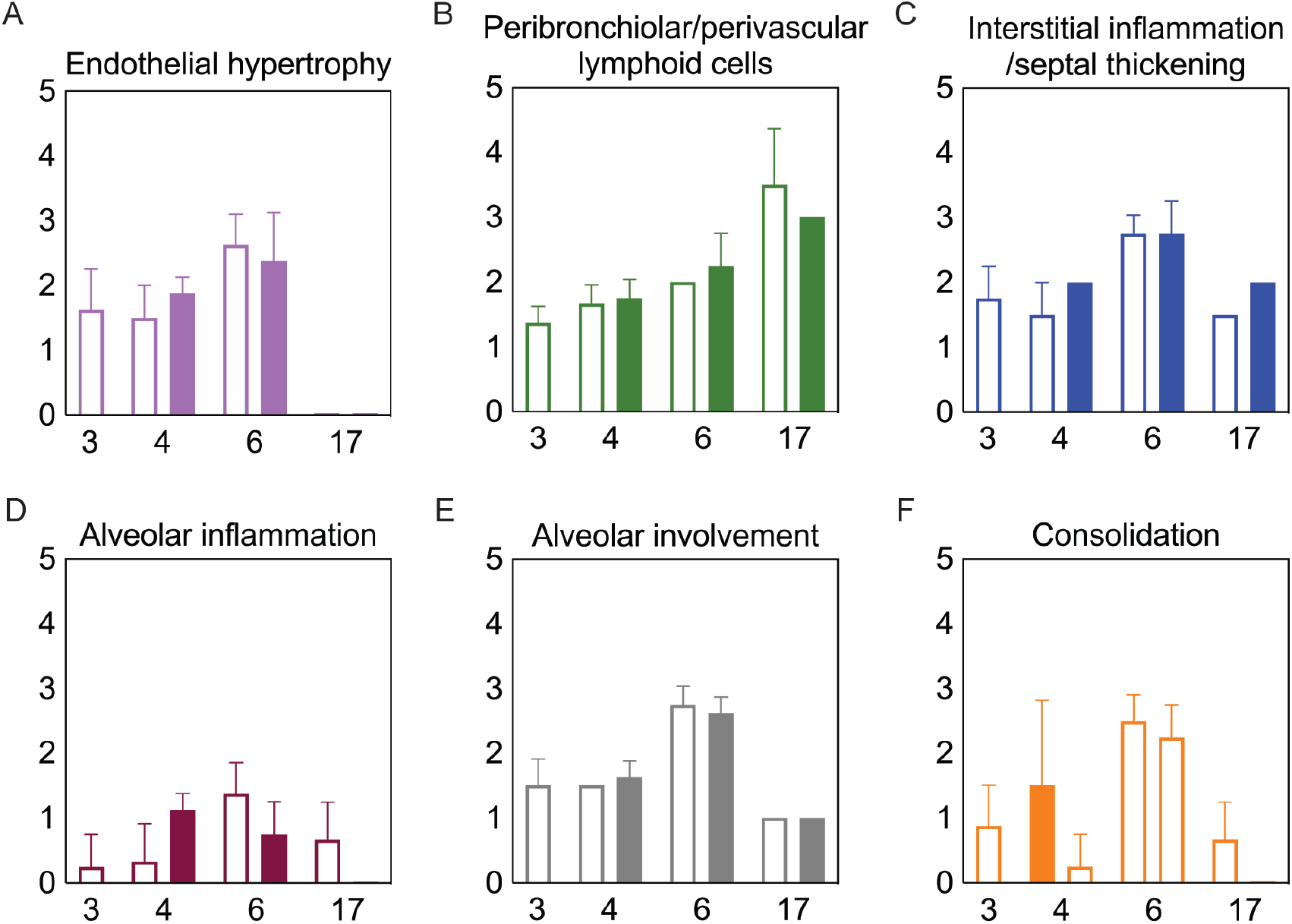
Lung pathology during SARS-CoV-2 infection and pneumococcal coinfection. Average endothelial hypertrophy (A), peribronchiolar/perivascular lymphoid cells (B), interstitial inflammation/septal thickening (C), alveolar inflammation (D), extent of alveolar involvement (E), and consolidation (F) in lungs of mice infected with SARS-CoV-2 (250 PFU; open bars) followed by 10^3^ CFU D39 at 3 or 5 d pvi (filled bars). Plots represent the mean ± standard deviation (SD) bars for combined male and female groups.

## Discussion

Currently, clinical data suggests variable, but moderate, frequency of bacterial coinfections in hospitalized COVID-19 patients^1–29^. The wide range of reported rates is, at least in part, due to heterogeneous study designs, variability in the disease severity, age, and/or comorbidities of each cohort, the collection and detection methods used, and the panel of pathogens screened. Further, the reduced transmission of many pathogens^104–108^ might have kept the rates of SARS-CoV-2-related bacterial pneumonia at an artificially low level during the COVID-19 pandemic. The results from this study suggest that we might expect more complications from bacterial pathogens going forward even in mild SARS-CoV-2 scenarios, which are becoming more common due to vaccine availability^109–111^.

Here, we used the K18-hACE2 mouse model to establish that SARS-CoV-2 infection increases the risk of bacterial coinfection in a time-dependent manner with increased disease severity, pulmonary bacterial burden, bacteremia, and neutrophilia. This time dependency is similar to that of influenza-bacterial coinfections, but the lethality during the SARS-CoV-2-pneumococcal coinfection (Figure 1) was delayed comparatively^77^ and some animals survived. In contrast, influenza-pneumococcal coinfections at similar doses consistently result in 100% lethality within 1-3 d pbi^77^. Although further studies are needed to assess the potential for more severe coinfections at later time points, this may indicate a larger window for administration of antibacterial therapies in coinfected patients.

Mechanisms that contribute to increased risk and severity of bacterial coinfection during acute pulmonary diseases are complex and varied^Reviewed in 36,39–41,45,84,85,112^. While the mechanisms for SARS-CoV-2-bacterial coinfections remain unknown, the similar time-dependent susceptibility during influenza may yield insight. We and others have shown that viral-induced changes to the number^67,69,70^ or functionality^70,72,113–115^ of AMΦs, which may be mediated by IFN-γ^55,115,116^, render these cells less capable of clearing bacteria. Here, SARS-CoV-2-pneumococcal coinfection did coincide with a virally induced reduction in AMΦ (Figure 3), which may suggest a contribution of these cells to the acquisition of bacteria during COVID-19 particularly when paired with evidence of a dysfunctional myeloid response in patients with severe infections^75^. Further studies to determine how a productive SARS-CoV-2 infection of AMΦ alters infection dynamics, their production of IFN, and their phagocytic capacity^86–89^ are needed. In addition, IFN-independent mechanisms of macrophage dysfunction should also be investigated because some studies suggest that RSV coinfection severity is mediated by Gas6/Axl polarization of AMΦ to non-antibacterial (M2) type cells^117^. Other mechanisms, including viral-mediated changes in bacterial receptor expression and binding^77,118–121^ and the degradation of epithelial tight junction integrity^122,123^ may also promote bacterial adherence during IAV or RSV infections, and some evidence suggests that these also occur during SARS-CoV-2 infection^124–126^.

Several studies have found that neutrophil dysfunction contributes to pathogenicity of IAV-pneumococcal coinfection, and this seems to be mediated by bacterial metabolism^82^ and type I IFNs^71,83,127^. However, unlike IAV-pneumococcal coinfections, type I IFNs were unchanged after SARS-CoV-2-pneumococcal coinfection (Figure S5) and neutrophil infiltration was only observed in coinfection at 7 d pvi (Figure 3A), suggesting that there may be different mechanisms underlying the enhanced pathogenicity of SARS-CoV-2 pneumococcal coinfection. This may, in part, be related to the low dose used here, where some studies have found that the SARS-CoV-related alterations to the IFN and iMΦ responses occur during more severe infections^128^. It was intriguing to see here that cytokine production was largely unchanged at 24 h pbi (Figure 4 and Figure S5), which is in contrast with the robust proinflammatory cytokine/chemokine production during other viral-bacterial coinfections^39–41,45,84,85^. Perhaps unexpectedly, several cytokines associated with severe COVID-19 and damaging cytokine overproduction (IL-6, IL-15, and IL-18)^129,130^ were reduced following coinfection at 5 d pvi (Figure 4).

Although coinfections are typically thought to be hyperinflammatory with enhanced disease severity, tissue inflammation does not seem to be altered during SARS-CoV-2-pneumococcal (Figure 5) or influenza-pneumococcal^55^ coinfections even with large neutrophil infiltrations^55,82^ (Figure 3A), at least within the first few days of coinfection. This may be owed to the nonlinearities between host immune responses, tissue inflammation, and disease severity^55,56^. Although the pathogenicity was increased during the coinfections at 5 d and 7 d pvi, there seemed to be little contribution from SARS-CoV-2, where the burden and distribution did not change within the first 24 h pbi (Figure 2) despite reduced CD8^+^ T cells in some groups (Figure 3G). In IAV-pneumococcal coinfections, invading bacteria result in robustly increased viral loads^55,68,82,131–133^ regardless of timing^55^ and viral dissemination in the lung is increased by 30-50%^55^. Our prior work^55^ suggests this is due to a combination of direct viral-bacterial interactions^97^ that lead to viral access to new areas of the lung in addition to increased virus production rates^68^ that may be mediated by alterations to the antiviral IFN response^98^. The lack of detection of SARS-CoV-2 in new areas of the lung may suggest that SARS-CoV-2 cannot as readily attach to pneumococcus like other viruses^97,134^, which is positive news given that pneumococci easily invade the blood^Reviewed in 135^ and SARS-CoV-2 affects numerous other organs^51–54^.

Although the long-term effects of viral-bacterial coinfections are not well studied, these data suggest they may be important where the SARS-CoV-2-bacterial coinfection resulted in lasting immunologic changes in recovered individuals. The higher macrophages and T cells (Figure 3) and their associated cytokines (Figure 4 and Figure S5) at 17 d pvi in animals recovered from bacterial coinfection is intriguing and suggests sustained immunopathology^55,56,136,137^. Many of the elevated responses are indicators of acute respiratory distress syndrome (ARDS)^138,139^ and are upregulated to promote tissue recovery and reduce pathology^140,141,142,143^. This was reflected in the slightly greater interstitial inflammation 17 d pvi (Figure 5) in coinfected animals. Interestingly, several cytokines were lower in animals that had recovered from bacterial coinfection with some below that of a naïve animal (Figures S5 and S6), which may also support a remodeling environment.

In summary, we used the transgenic K18-hACE mouse model^144^ to establish that a low dose SARS-CoV-2 infection increases the risk of pneumococcal coinfection in a time-dependent manner. The data importantly highlight many differences with other viral-bacterial coinfections and the need for further studies to clarify the host-pathogen interplay that enhance susceptibility and pathogenicity during SARS-CoV-2-bacterial coinfection. This information may be crucial going forward, particularly because a sustained immune activation following coinfection suggests an increased risk of developing ARDS even in patients with mild COVID-19. In addition, as new SARS-CoV-2 variants emerge and nonpharmaceutical measures, such as wearing masks and physical distancing, become less common, we might anticipate an increase in risk of bacterial transmission and acquisition in COVID-19-infected individuals.

## Materials and Methods

### Mice

Adult (10-13 week old) male and female K18-hACE2 transgenic mice (B6.Cg-Tg(K18-ACE2)2Prlmn/J) were obtained from Jackson Laboratories (Bar Harbor, ME). Mice were housed in groups of 4 in solid–bottom polysulfone individually ventilated cages (Allentown BCU) in rooms maintained on a 12:12-hour light:dark cycle at 22 ± 2°C with 30-70% humidity in the Regional Biocontainment Laboratory (animal biosafety level 3 facility) at UTHSC (Memphis, TN). Mice were acclimated for 1 day before being lightly anesthetized with 2% inhaled isoflurane (Baxter, Deerfield, IL) and implanted subcutaneously with an IPTT300 transponder (Bio Medic Data Systems, Seaford, DE) for identification and temperature monitoring, followed by an additional 3 days of acclimation before inclusion in the experiments. Envigo irradiated rodent diet (catalog no. 7912) and autoclaved water were available ad libitum during the acclimation and study periods; gel food and hydrogel were provided at the time of infection. All experimental procedures were performed under protocol 20-0132 approved by the Animal Care and Use Committee at University of Tennessee Health Science Center (UTHSC) under relevant institutional and American Veterinary Medical Association (AVMA) guidelines and were performed in a biosafety level 3 facility that is accredited by the American Association for Laboratory Animal Science (AALAS).

### Infection experiments

All experiments were done using 2019-nCoV/USA-WA1/2020 (BEI Resources NR-52281) (SARS-CoV-2) and type 2 pneumococcal strain D39. The viral infectious dose [plaque forming units (PFU)] was determined by plaque assay of serial dilutions on Vero E6 cells. Virus seed stocks were sequenced using next-generation sequencing with ARTIC primers on the Illumina MiSeq. Bacterial infectious dose [colony forming units (CFU)] was determined by using serial dilutions on tryptic soy agar plates supplemented with 3% sheep erythrocytes (TSA). Doses of virus and bacteria were selected that elicited mild-moderate disease independently to ensure that changes in disease severity following coinfection would be evident. Frozen stocks were diluted in sterile PBS and administered intranasally to groups of 4 mice, lightly anesthetized with 2.5% inhaled isoflurane (Baxter, Deerfield, IL) in a total volume of 50 μl (25 μl per nostril). Mice were inoculated with either PBS or SARS-CoV-2 at day 0 then with 10^3^ CFU of D39 or PBS, either 3 or 5 days later. Weight loss, temperature change, appearance, respiratory effort, behavior, and dehydration were scored (scale 1 to 3) at the onset of infection and each subsequent day to monitor illness and mortality. Mice were euthanized if they lost 30% of their starting body weight or became moribund based on clinical scores (single category score of 3 or cumulative score of ≥9 in respiratory distress, dehydration, temperature reduction, behavior/mobility, body condition/appearance).

### Harvest and processing of lungs and blood

Mice were euthanized by 33% isoflurane inhalation. Lungs were aseptically harvested, washed in PBS, and fixed in 10% neutral buffered formalin for histology or digested with collagenase (1 mg/ml, Sigma C0130) and physical homogenization against a 40 μm cell strainer for immune cell staining. Lung digest supernatants were used to quantify cytokines and chemokines and to determine viral and bacterial titers as above; bacterial titers were also measured in peripheral blood. Following red blood cell lysis, lung cells were washed in staining buffer (PBS, 5mM EDTA, 10mM HEPES, and 0.5% bovine serum albumin), counted with trypan blue exclusion using a Cell Countess System (Invitrogen, Grand Island, NY), and prepared for flow cytometric analysis as described below.

### Flow cytometric analysis

Flow cytometry (BD FACSAria; San Jose, CA) was performed on single cell suspensions after Fc receptor blocking (TruStainFcX, Biolegend) and viability staining (Zombie Violet Fixable Viability, Biolegend), 25 min surface staining, and fixation (BD Cytofix). The followed anti-mouse antibody panels were used for cell subset analysis: CD45 (clone 30-F11, Pe-Cy7, Biolegend), CD3e (clone 145-2C11, FITC, Biolegend), CD4 (clone RM4-5, V500, BD Biosciences), CD8α (clone 53-6.7, PerCP-Cy5.5, Biolegend), CD19 (clone 6D5, PE, Biolegend), CD335 (clone 29A1.4, APC-Fire750, Biolegend), and CD69 (clone H1.2F3, APC, Biolegend) or CD45 (clone 30-F11, Pe-Cy7, Biolegend), Ly6G (clone 1A8, PerCP-Cy5.5, Biolegend), F4/80 (clone BM8, PE, eBioscience), CD11b (clone M1/70, V500, BD Biosciences), CD11c (clone N418, APC-Fire750, Biolegend), MHC-II (clone I-A/I-E, FITC, eBioscience), and CD69 (clone H1.2F3, APC, Biolegend). The data were analyzed using FlowJo 10.7.2 (Tree Star, Ashland, OR). Data were cleaned using the flowAI application^145^ followed by gating viable cells from a forward scatter/side scatter plot, singlet inclusion, and viability dye exclusion. CD45^+^ cells were selected for further analyses. Neutrophils (Ly6G^hi^), alveolar macrophages (AMΦ) (F4/80^hi^CD11c^hi^CD11b^-^MHC-II^low/-^), inflammatory/exudate macrophages (iMΦ) (F4/80^hi^CD11c^hi^CD11b^+^MHC-II^mid/hi^), other monocyte/macrophage populations (F4/80^mId^CD11c^mid^CD11b^+^ and F4/80^mid^CD11c^-^CD11b^+/-^), NK Cells (CD3e^-^CD19^-^ CD335^+^), CD4 T cells (CD3^+^CD8^-^CD4^+^CD335^-^), CD8 T cells (CD3^+^CD8^+^CD4^-^CD335^-^), NK-T cells (CD3e^+^CD335^+^), B cells (CD3e^-^CD19^+^), and recently activated subsets thereof (CD69^+^) were gated as in Fig S2.

### Cytokine and chemokine quantification

Cytokines G-CSF (CSF-3), GM-CSF, IFN-γ, IL-1α, IL-1β, IL-2, IL-3, IL-4, IL-5, IL-6, IL-9, IL-10, IL-12p70, IL-13, IL-15/IL-15R, IL-17A (CTLA-8), IL-18, IL-22, IL-23, IL-27, IL-28, IL-31, LIF, MCP-3 (CCL7), M-CSF, TNF-α) and chemokines (ENA-78 (CXCL5), eotaxin (CCL11), GROα (CXCL1), IP-10 (CXCL10), MCP-1 (CCL2), MIP-1α (CCL3), MIP-1β (CCL4), MIP-2α (CXCL2), RANTES (CCL5) were measured in lung supernatant by Luminex and ELISA (IFN-α,β). Before use, cell debris and aggregates were removed by centrifugation at 4°C, 400 x *g*. ProcartaPlex magnetic bead cytokine/chemokine plates (Invitrogen) were prepared according to the manufacturer’s instructions. Data were acquired using a MagPix (Luminex) with Luminex xPonent software (v4.2) and analyzed with the ProcartaPlex Analysis App (ThermoFisher Connect). ELISAs for IFNa and IFNβ (PBL Assay Science) were prepared according to the manufacturer’s instructions, read at 450 nm, and analyzed using GraphPad Prism 9.2.0. Mean concentrations of duplicate samples were calculated by the construction of standard curves using a weighted 5PL and 4PL regression for the ProcartaPlex and ELISA data, respectively. Absolute quantities of each cytokine/chemokine were calculated based on the mean concentration of replicate samples normalized to the lung supernatant volume collected during tissue processing. Internal plate controls were used to adjust values obtained between plates and fold changes in cytokine and chemokine quantities were calculated for each animal, normalized to the average of naïve controls (pooled males/females).

### Histology

Following euthanasia and tissue removal as above, lungs were continually fixed in 10% neutral-buffered formalin solution (NBF; ThermoFisher Scientific, Waltham, MA) before being embedded in paraffin, sectioned at 4μm, and mounted on positively charged glass slides (Superfrost Plus; Thermo Fisher Scientific, Waltham, MA). Tissue sections were stained with hematoxylin and eosin (H&E) or subjected to immunohistochemical (IHC) staining to detect SARS-CoV-2 antigen. Tissue sections were deparaffinized and rehydrated before undergoing antigen retrieval in a citrate-based solution (pH 6.0) (Vector Laboratories, Burlingame, CA) at 97°C. For IHC, a primary monoclonal antibody against SARS-CoV-2 nucleoprotein (NP) (Sino Biological, Wayne, PA) was used at 1:1000 followed by a biotinylated anti-rabbit antibody (Vector Laboratories, Burlingame, CA) at 1:200, the Vectastain Elite ABC-HRP kit (Vector Laboratories, Burlingame, CA), and 3,3’-Diaminobenzidine (DAB) solution development. Stained sections were counterstained with hematoxylin, dehydrated, and examined by a pathologist blinded to the experimental group assignments. To quantify the extent of viral infection in the lungs, digital images of whole lung sections stained for viral antigen were first captured using the Aperio ScanScope XT Slide Scanner (Aperio Technologies, Inc., Vista, CA). The areas of both the entire lung parenchyma (alveoli and bronchioles) and the virus-positive regions were outlined manually with areas determined using ImageScope software (Aperio Technologies, Inc.). Representative images and quantitative analyses of viral spread and lung pathology during infection are shown in Figure 2 and Figure 5, respectively.

### Statistical Analysis

Significant differences in Kaplan-Meier survival curves were calculated using the log-rank test. Linear values of lung and blood bacterial loads, viral loads, immune cells, cytokines/chemokines were compared using unpaired t-tests with Welch correction or Mann-Whitney test where appropriate (GraphPad Prism 9.2.0 and Rv4.0.3). The confidence interval of significance was set to 95%, and *P* ≤ 0.05 was considered significant.

## Data availability

The following reagent was deposited by the Centers for Disease Control and Prevention and obtained through BEI Resources, NIAID, NIH: SARS-Related Coronavirus 2, Isolate USA-WA1/2020, NR-52281.

## Acknowledgments

We thank the staff of the Regional Biocontainment Laboratory and Deidre Daria, Ph.D. for technical support, and Jyothi Parvathareddy and Dong Yang for the generation and characterization of viral stocks.

This work was supported by the Institute for the Study of Host Pathogen Systems at UTHSC and NIH grant number AI139088.

