## Supplemental Material for "Time-Dependent Increase in Susceptibility and Severity of Secondary Bacterial Infection during SARS-CoV-2 Infection"

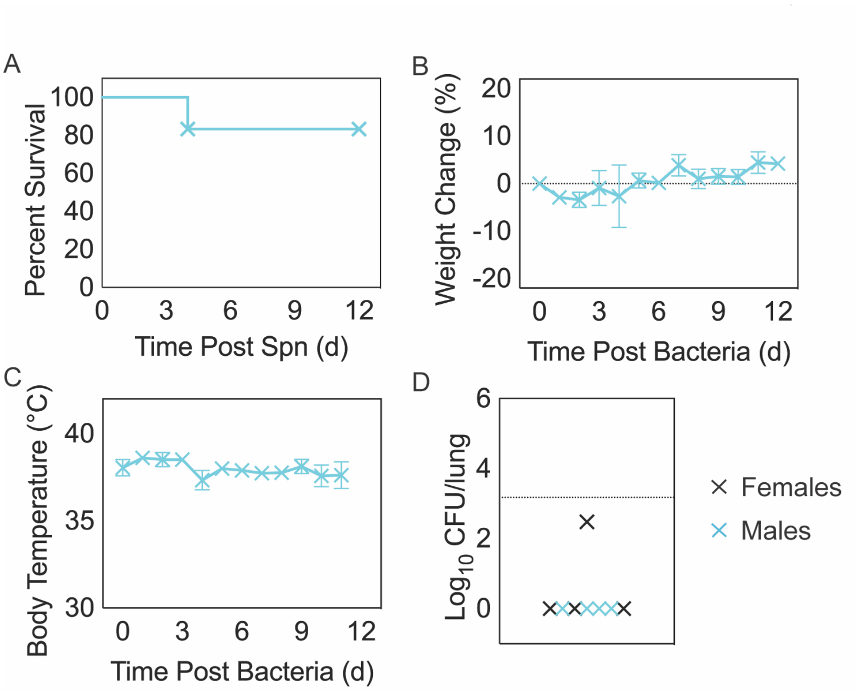


**Figure S1: *Streptococcus pneumoniae* infection in naïve K18-hACE2 mice.** Kaplan-Meier survival curve (A), percent weight loss (B), and temperature (C) of mice infected with $10^3$ CFU D39. Data are shown as the mean ± standard deviation (SD). Lung bacterial loads (CFU/lung) (D) in female (black) and male (cyan) mice infected with $10^3$ CFU D39 for 24 hours.

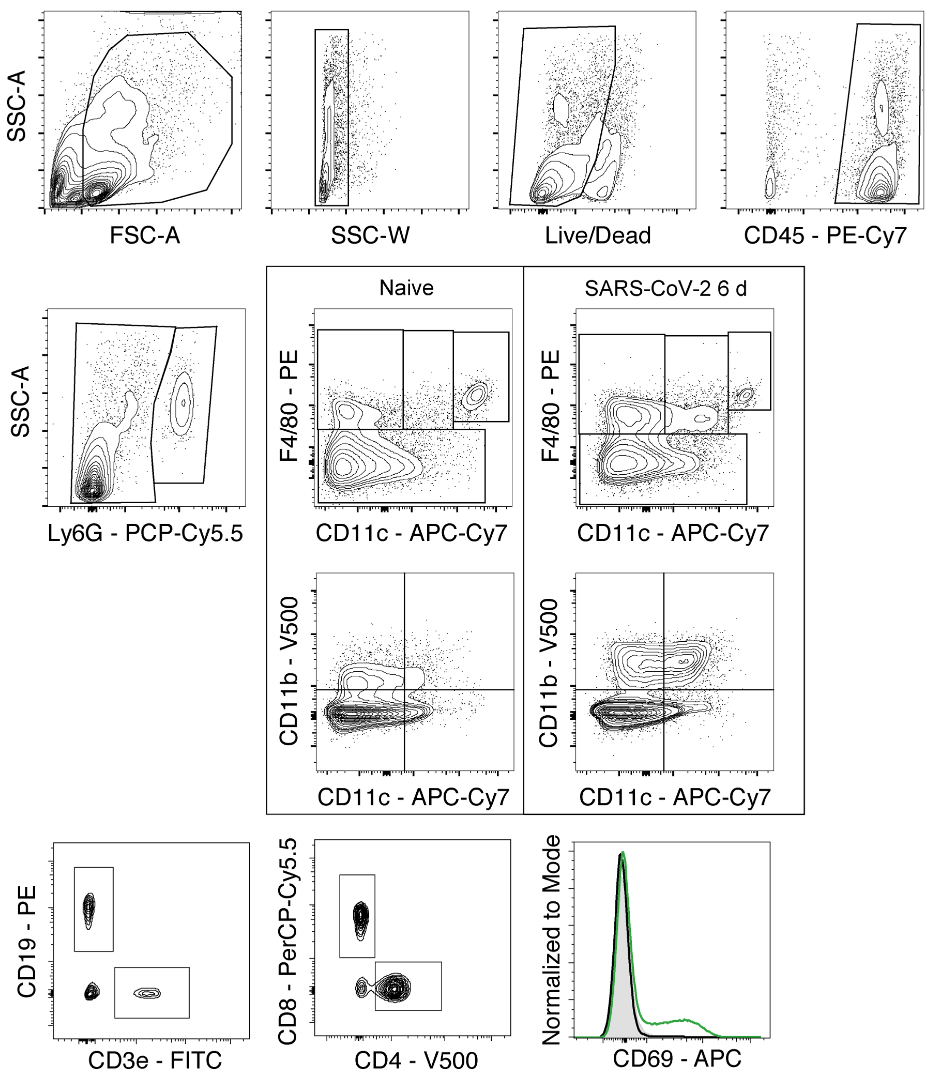


**Figure S2: Flow cytometry gating scheme for lung cell analysis.** Viable immune cells were first gated on forward scatter (FSC-A) and side scatter (SSC-A), as singlets, fixable viability dye negative, and CD45$^{+}$ (top row). Neutrophils (Ly6G$^{hi}$) were then gated and excluded from remaining parent populations. Monocyte and macrophage (MΦ) subsets were gated based on expression of CD11c and F4/80 with alveolar macrophages (AMΦ) sub-gated as F4/80$^{hi}$CD11c$^{hi}$CD11b$^{-}$MHC-II$^{low/-}$, inflammatory macrophages (iMΦ) as F4/80$^{hi}$CD11c$^{hi}$CD11b$^{+}$MHC-II$^{mid/hi}$, and additional subsets as F4/80$^{mid}$CD11c$^{mid}$CD11b$^{+}$ and F4/80$^{mid}$CD11c$^{-}$CD11b$^{+/-}$. Following MΦ exclusion, B cells were gated as CD3e$^{-}$CD19$^{+}$ and T cells were gated as CD3e$^{+}$ and subgated into CD8$^{+}$ T cells (CD3e$^{+}$CD8$^{+}$CD4$^{-}$CD335$^{-}$), CD4$^{+}$ T cells (CD3e$^{+}$CD8$^{-}$CD4$^{+}$CD335$^{-}$), and NK T cells (CD3e$^{+}$CD335$^{+}$). Natural killer (NK) cells were gated CD3e$^{-}$CD19$^{-}$CD335$^{+}$. Surface CD69 expression was used to quantify activation in all gated populations.

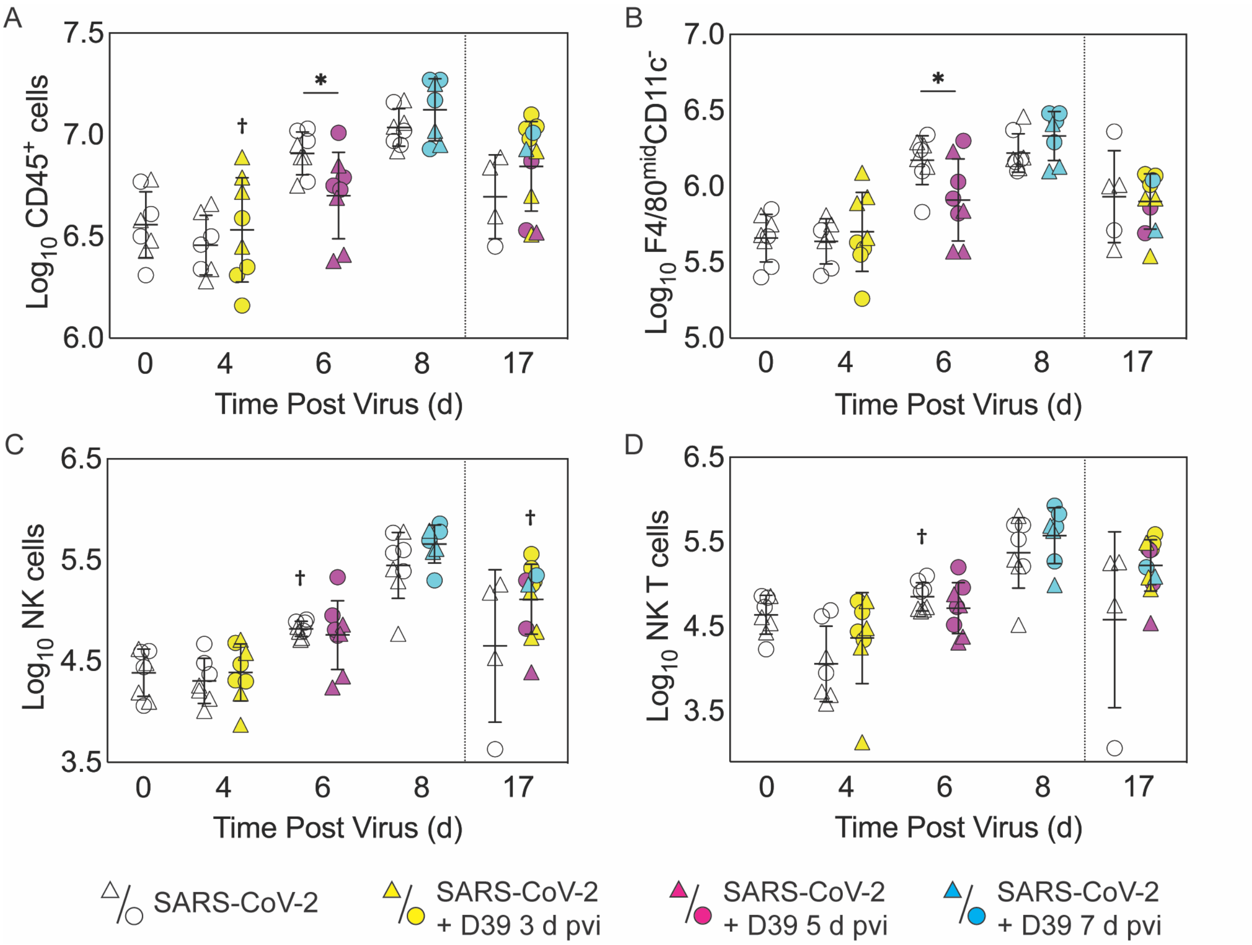


**Figure S3: Quantification of additional immune cell populations.** Total $CD45^+$ cells (A), $F4/80^{mid}CD11c^-CD11b^{+/-}$ cells (B), NK cells (C), and NK-T cells (D) in the lungs of female (circles) and male (triangle) mice infected with SARS-CoV-2 (250 PFU; open symbols) followed by infection with $10^3$ CFU D39 at 3 d (yellow), 5 d (magenta), or 7 d (cyan) pvi. Each symbol represents a single mouse and the mean ± standard deviation (SD) are for combined male and female groups. Significant differences are indicated by *, $P < .05$ for comparisons between indicated groups and as †, $P < .05$ for differences between males and females within a group or between coinfection times within 17 d group. Plots depicting additional cells are shown in Figure 3 of the main text and the flow cytometry gating scheme is in Figure S2.

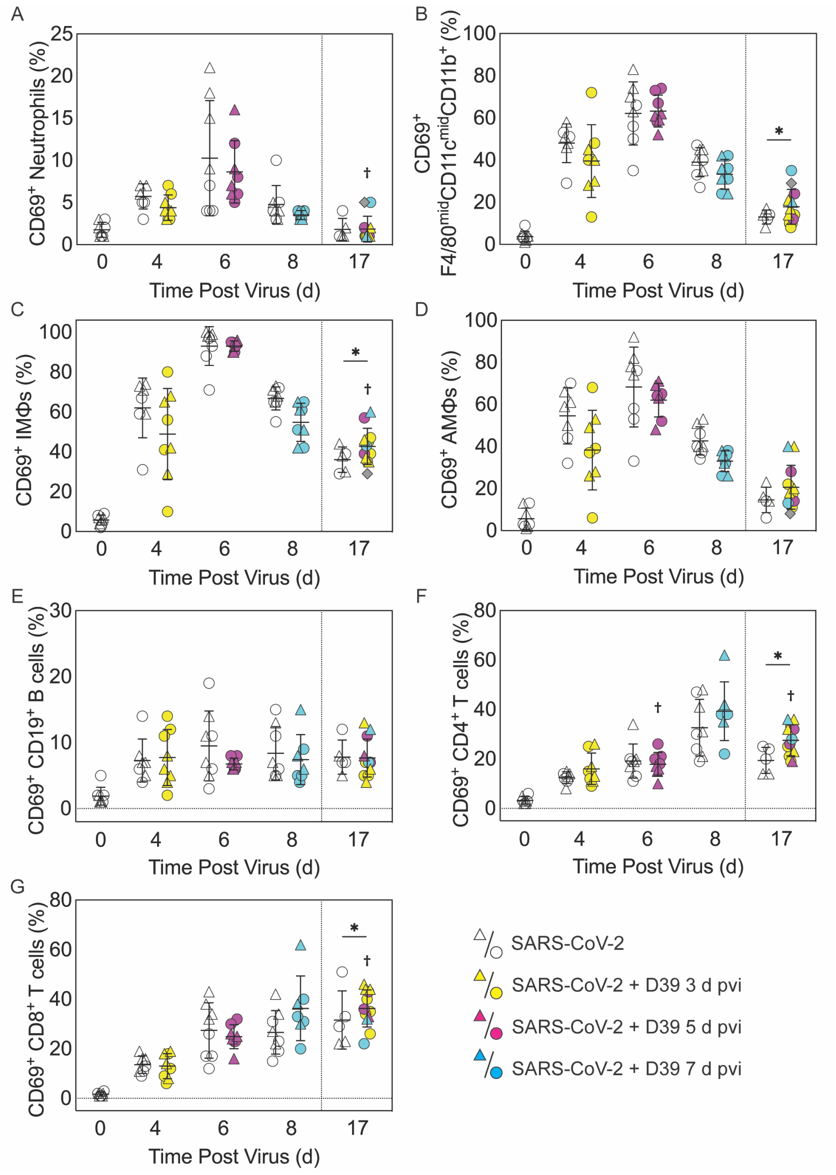


**Figure S4: Immune cell activation during SARS-CoV-2 infection and pneumococcal coinfection.** The percentage of $CD69^+$ neutrophils (A), $F4/80^{mid}CD11c^{mid}CD11b^+$ (B), iMΦ ($F4/80^{hi}CD11c^{hi}CD11b^+MHC\text{-}II^{mid/hi}$) (C) AMΦ ($F4/80^{hi}CD11c^{hi}CD11b^-MHC\text{-}II^{low/-}$) (D), $CD19^+$ B cells (E), $CD4^+$ T cells (F), and $CD8^+$ T cells (G) in the lungs of female (circles) and male (triangle) mice infected with SARS-CoV-2 (250 PFU; open symbols) followed by infection with $10^3$ CFU D39 at 3 d (yellow), 5 d (magenta), or 7 d (cyan) pvi. Each symbol represents a single mouse and the mean ± standard deviation (SD) are for combined male and female groups. Significance comparisons were done using the absolute number of cells per lung and differences are indicated by *, *P* < .05 for comparisons between indicated groups and as †, *P* < .05 for differences between males and females within a group or between coinfection times within 17 d group.

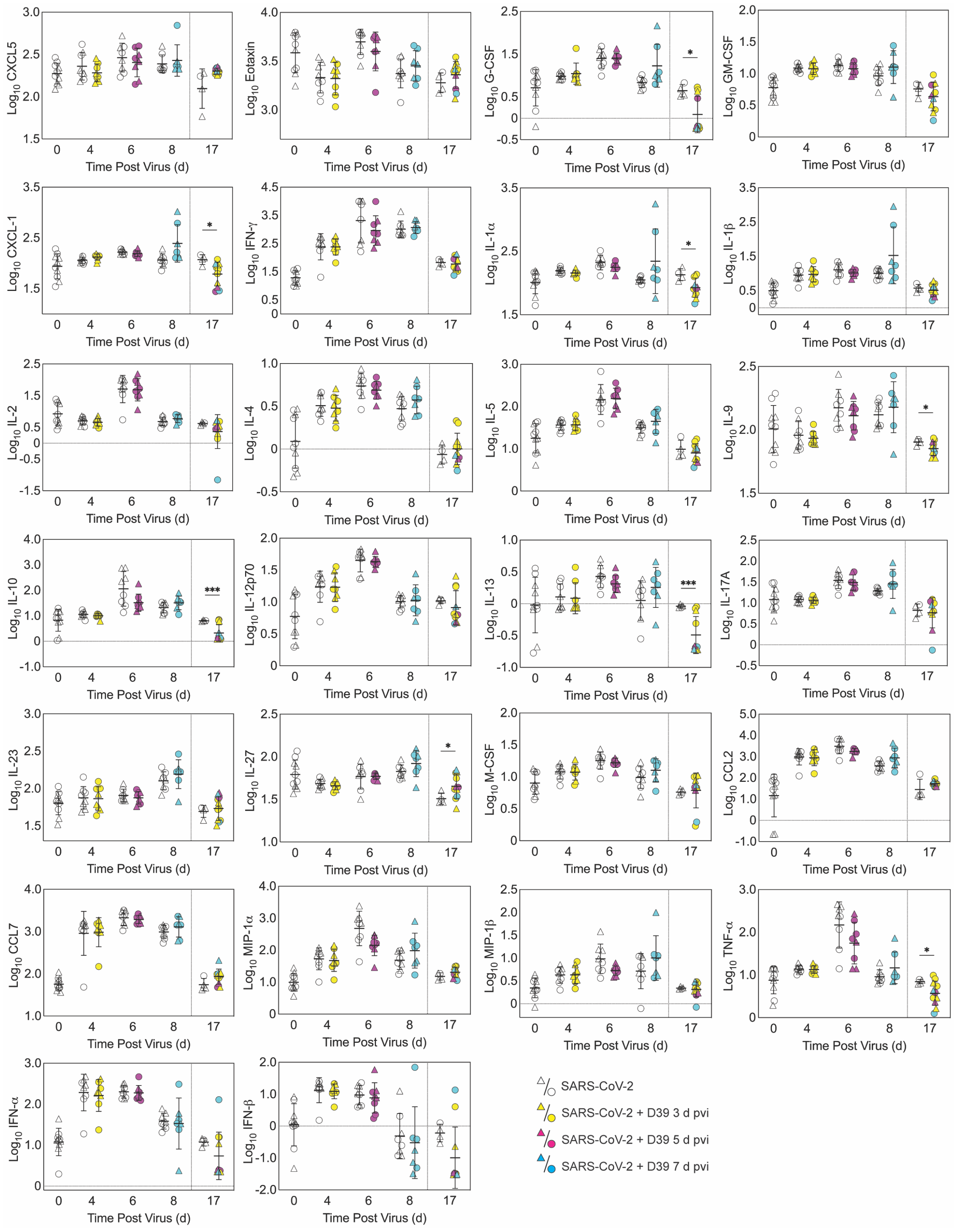


**Figure S5: Absolute quantities of pulmonary cytokines and chemokines during SARS-CoV-2 infection and pneumococcal coinfection.** The absolute picograms ($\log_{10}$) of cytokines and chemokines in the lungs of female (circles) and male (triangle) mice infected with SARS-CoV-2 (250 PFU; open symbols) followed by infection with $10^3$ CFU D39 at 3 d (yellow), 5 d (magenta), or 7 d (cyan) pvi. Each symbol represents a single mouse and the mean ± standard deviation (SD) are for combined male and female groups. Significant differences are indicated by *, $P < .05$; **, $P < .01$; and ***, $P < .001$ for comparisons between indicated groups. Plots depicting additional cytokine and chemokine quantities (absolute $\log_{10}$ picograms) are in Figure 4 of the main text and a heatmap representing the normalized quantity (average $\log_2$ change over naïve) is in Figure S6.

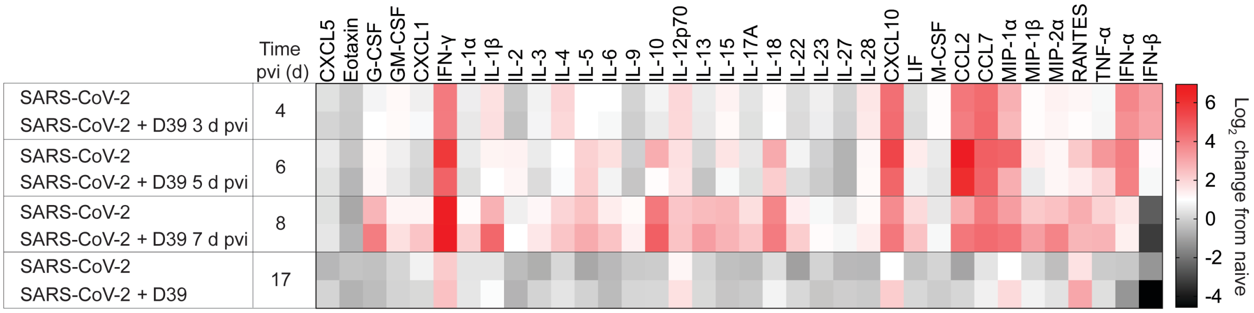


**Figure S6: Fold change over naïve of pulmonary cytokines and chemokines during SARS-CoV-2 infection and SARS-CoV-2-pneumococcal coinfection.** Heatmap representing the normalized quantity (average $\log_2$ change over naïve) of 36 cytokines and chemokines in the lungs of mice infected with SARS-CoV-2 (250 PFU) followed by $10^3$ CFU D39 at 3, 5, or 7 d pvi. Plots depicting absolute $\log_{10}$ picograms (pg) of measured cytokines and chemokines are in Figure 4 and Figure S5.
